# A Data-Driven Optimization Method for Coarse-Graining Gene Regulatory Networks

**DOI:** 10.1101/2022.08.10.503498

**Authors:** Cristian Caranica, Mingyang Lu

## Abstract

One major challenge in systems biology is to understand how various genes in a gene regulatory network (GRN) collectively perform their functions and control network dynamics. This task becomes extremely hard to tackle in the case of large networks with hundreds of genes and edges, many of which have redundant regulatory roles and functions. The existing methods for model reduction usually require the detailed mathematical description of dynamical systems and their corresponding kinetic parameters, which are often not available. Here, we present a data-driven method for coarse-graining large GRNs, named SacoGraci, using ensemble-based mathematical modeling, dimensionality reduction and gene circuit optimization by Markov Chain Monte Carlo methods. SacoGraci requires network topology as the only input and is robust against errors in GRNs. We benchmark and demonstrate its usage with synthetic, literature-based, and bioinformatics-derived GRNs. We hope SacoGraci will enhance our ability to model the gene regulation of complex biological systems.

## Introduction

One of the major challenges in systems biology is to understand how gene regulatory networks (GRNs) enable the creation and maintenance of cellular states and how they control and drive cellular state transitions in biological processes, such as cell differentiation and disease progression (Gerstein et al., 2012; Schwikowski et al., 2000).Common ways to construct GRNs (Katebi et al., 2021; Shahzad and Loor, 2012) are either by the bottom-up approach (Feist et al., 2009; Kulkarni et al., 2012; Thiele and Palsson, 2010), in which gene regulatory interactions are derived from literature data, or by the top-down approach (Pratapa et al., 2020) in which bioinformatics methods are applied on genome-wide omics data (Aibar et al., 2017; Deshpande et al., 2022; Margolin et al., 2006; Matsumoto et al., 2017; Pranzatelli et al., 2018), such as transcriptomics data. The constructed GRNs often contain a large number of genes and edges, making it hard to understand how GRNs operate and control their dynamics.

One strategy to address these issues is network coarse-graining, which constructs small core regulatory circuits that capture the dynamical behavior of large GRNs. Large GRNs often contain many genes and edges with redundant regulatory roles and functions, thus the redundant components can be combined into simple network models (Chauhan et al., 2021; Lu et al., 2013; Tripathi et al., 2022). Compared to a large GRN, a small circuit model is more likely to capture the most important network functions and to reveal the roles of genes and the relationship between genes. Moreover, it is easier to fix errors or make changes in small GRNs than in those large counterparts because of much reduced searching space.

Coarse-graining, sometimes referred to as model reduction, is a popular technique in physics, chemical, and engineering disciplines (Atilgan et al., 2001; Ingólfsson et al., 2014; Kmiecik et al., 2016; Lu and Ma, 2005; Lu et al., 2006; VanWart et al., 2012). Various coarse-graining methods have been developed in systems biology to model GRNs (Erban et al., 2006; Sinitsyn et al., 2009; Snowden et al., 2017). The most representative ones are based on timescale exploitation (Prescott and Papachristodoulou, 2014), optimization (Danø et al., 2006; Maurya et al., 2009), lumping (Dokoumetzidis and Aarons, 2009) or singular value decomposition (Meyer-Bäse and Theis, 2008). However, most existing methods require the knowledge of the mathematical rate equations and detailed kinetic parameters of the full models, which is not available in many cases. Moreover, it is often difficult for the existing methods to establish high quality models from GRNs containing missing or inaccurate regulatory interactions.

To alleviate these limitations, here we present a new GRN coarse-graining method, named *<u>Sa</u>mpling <u>co</u>arse-<u>Gr</u>ained <u>ci</u>rcuits* (SacoGraci), to construct optimized small circuit topologies that capture the gene expression states of a large GRN. SacoGraci first clusters the genes and models describing the network’s behavior, followed by an optimization process that samples coarse-grained circuits (CGCs) producing gene expression patterns similar to those of the GRN. As the core of SacoGraci, network optimization uses a new scoring function and multiple sampling schemes. Here, the scoring function quantifies the dissimilarity of gene expression states between the small circuit and the large GRN based on the simulations of gene expression by an ensemble-based mathematical modeling method named RACIPE (Huang et al., 2017; Kohar and Lu, 2018). Because RACIPE captures GRN’s gene expression states from an ensemble of models with randomly generated kinetic parameters (Huang et al., 2018, 2020; Katebi et al., 2020; Ramirez et al., 2020; Su et al., 2022), there is no need to specify or optimize kinetic parameters in each score evaluation. Such a feature allows us to focus on the optimization of circuit topology by a Markov Chain Monte Carlo (MCMC)-based sampling scheme (Landau and Binder, 2009), where the sampling space is determined by the topology of the large GRN. SacoGraci has the advantage over most existing methods in that it only requires the GRN topology as the input; it is robust against errors in large GRNs; and the optimization is independent to the choice of model parameters.

In the following, we will first provide an overview of SacoGraci, with the detailed algorithms explained in the Methods section and Supporting Information. We will then show the benchmark of SacoGraci on a large series of GRNs expanded from two synthetic gene circuits and perturbed in the network topology by different levels. Finally, we will illustrate its applications on four biological GRNs, including both literature-based and bioinformatics-derived networks.

## Results

### Overview of the coarse-graining algorithm SacoGraci

SacoGraci takes the topology of a large gene regulatory network (GRN) as the input and identifies the optimal coarse-grained circuits (CGCs) that best capture the gene expression states of the full GRN. The workflow of SacoGraci is illustrated in **Fig.1**. First, RACIPE is applied to the full GRN to simulate the stable steady-states gene expression profiles of an ensemble of mathematical models with randomly selected kinetic parameters. Second, clustering analysis is applied to the simulated gene expression profiles to identify model clusters, each of which defines a *network state*, and gene clusters, each of which defines a node in the CGC. The number of network states and the number of circuit nodes can be either determined by biological context or be considered as hyperparameters for optimization. Third, an optimization process is then applied to sample candidate circuits and obtain the optimal CGCs by a Markov Chain Monte Carlo (MCMC) method using a scoring function that quantifies the mismatch between the network states of the CGC and those of the full GRN. The regulatory interactions are sampled according to edge type (*i*.*e*., activation, inhibition, or no interaction) distributions determined from the full GRN. During the score evaluation, RACIPE is applied to each candidate circuit to obtain its stable steady-state gene expression profiles. Details of SacoGraci procedures are described in the Methods section.

**Figure 1.**
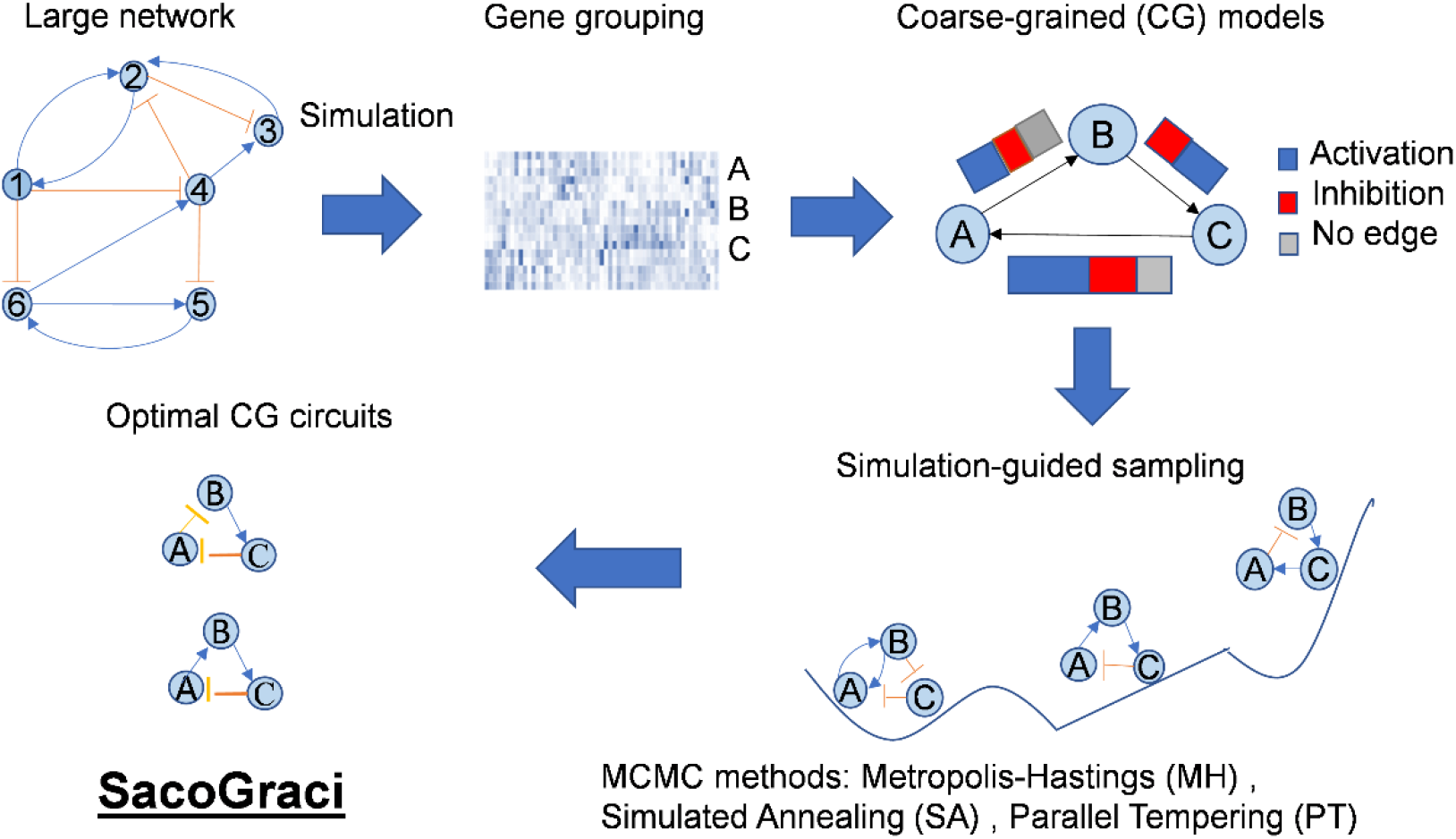
Workflow of the gene network coarse-graining algorithm. The goal of algorithm is to construct, from a large gene regulatory network, coarse-grained circuits (CGCs) models that capture the network’s behavior. First, given the topology of the full network, mathematical modeling method RACIPE simulates the steady-state gene expression profiles from an ensemble of randomly generated models. Second, cluster analysis is applied to simulated data to identify clusters of genes, which represent coarse-grained nodes, and clusters of gene expression profiles, which represent network states. Third, CGCs are constructed by Markov Chain Monte Carlo (MCMC)-based optimization, where circuit edges are chosen according to the regulatory interactions of the full network. The scoring function of the optimization is based on the similarity of the states of CGC and the states of the full network.

### Benchmarking SacoGraci using synthetic circuits

We first benchmarked SacoGraci in identifying high quality CGCs from a large GRN using two small synthetic circuits. We started from each small circuit and expanded it to a large network with redundant regulatory interactions derived from the small circuit. In the benchmark test, we applied SacoGraci to the large network (the input) and evaluated whether the identified CGC recovers not only the gene expression distribution of the original circuit, but also its topology (ground truth). Many biological networks may contain inconsistent edges (*i*.*e*., different excitatory or inhibitory edge types) even for genes with similar roles. Due to the availability of biological data on gene regulatory interactions, literature-derived GRNs may also contain inaccurate and missing edges. Thus, to evaluate the robustness of the network coarse-graining, we also tested SacoGraci on large networks with different levels of edge perturbations.

The first test case is a circuit of three coupled toggle-switches (diagram in **Fig.2A**). To generate the large network, we expanded each of the six nodes of the original circuit to four genes (details in Methods). The final large network contains 24 genes (diagram in **Fig.2B**), where genes with the same color correspond to the colored node of the original circuit. We clustered the models and the genes using hierarchical clustering analysis (HCA) with Ward.D2 (Ward, 1963) and one minus Pearson correlation as the distance function. As shown in the heatmaps of simulated gene expression profiles generated by RACIPE, the gene expression profiles from the large network (**Fig.2F**) form clusters of expression patterns similar to those obtained from the original circuit (**Fig.2E**). Genes corresponding to the same node were also clustered together, indicating that they play similar roles in network behavior. Thus, we chose four network states and four gene clusters for network coarse graining. The network states were clearly observed in the scatterplots of gene expressions projected onto the first two principal components (PCs) of large GRN’s simulated data. (**Fig.2HI**).

From the expanded network, perturbed networks for six different perturbation levels (each level with ten networks) were generated by randomly deleting, adding, and changing the signs of a proportion of edges, as described in Methods. These perturbed networks were used as the inputs to test SacoGraci (i.e., the perturbed network defines the edge type distributions for sampling CGCs). An example of a 30% edge-perturbed network is shown in **Fig.2C**. For this perturbation level, SacoGraci identified the CGC shown in **Fig.2D** as the best scored last-iteration circuit across all sampling methods. The optimized CGC generates similar network states (**Fig.2GJ**) as the large network. The CGC also nicely captures the network states of the original circuit (**Fig.2EH**), except for a lower proportion of models in the 3^rd^ state (blue) from the CGC.

**Figure 2.**
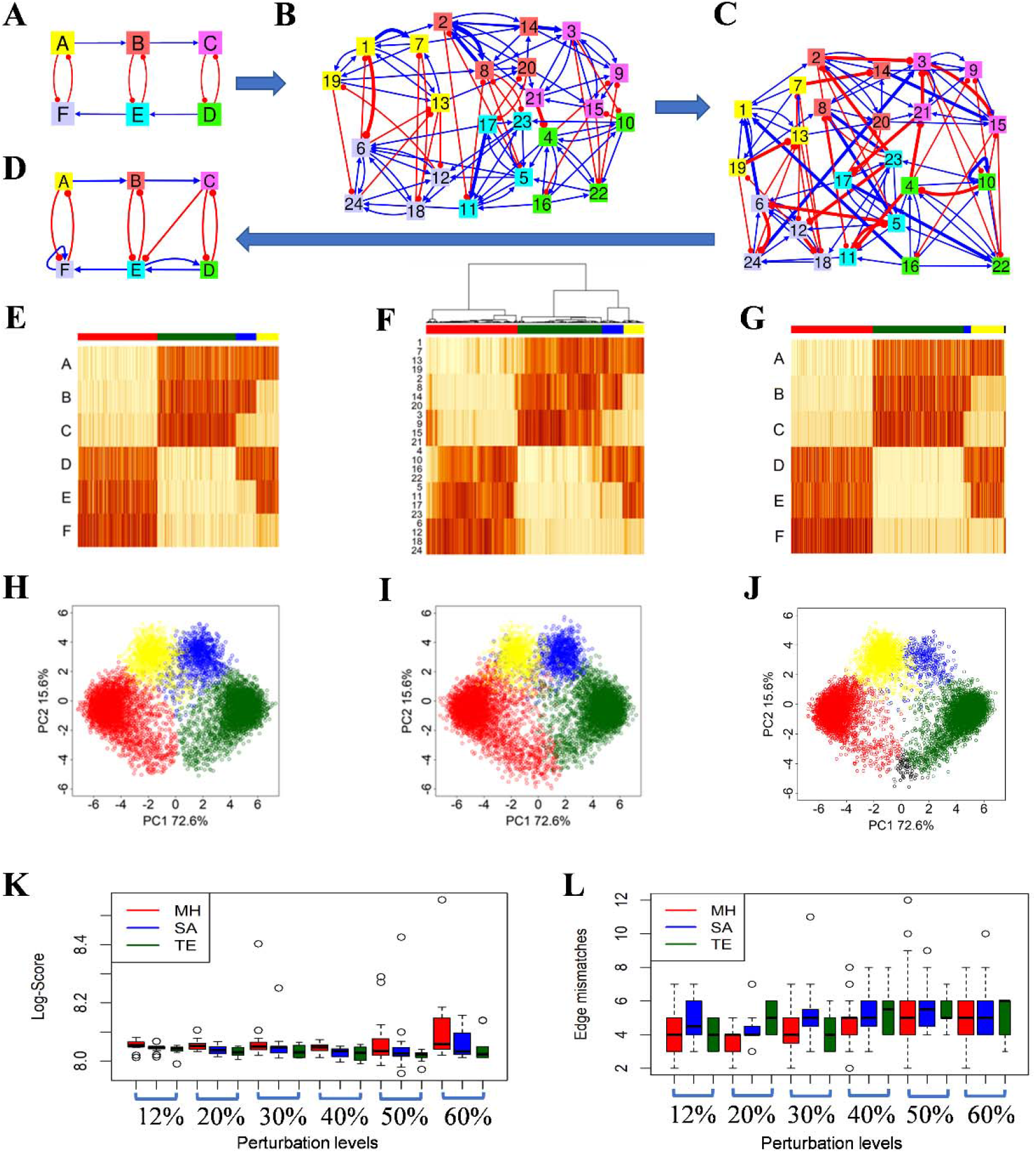
Benchmark tests on a synthetic gene circuit of three coupled toggle switches. **(A)** Topology of the original circuit. **(B)** Topology of an expanded network. Every node of the original circuit is expanded into four nodes of the same color. Edges that will not be present in the perturbed network are shown in bold. **(C)** Topology of a perturbed network with a 30% edge perturbation level. Edges that have been added to the expanded network or have changed the sign are shown in bold. **(D)** Topology of the best last-iteration CGC obtained after running all MCMC methods for networks with 30% perturbation level. (**E**) Heatmap of the RACIPE-simulated data for the original circuit. **(F)** Heatmap of simulated gene expression of the expanded network. It shows the hierarchical clustering analysis (HCA) of the RACIPE models using Ward.D2 as linkage method and 1 minus Pearson correlation as distance. Four well-separated clusters are determined based on HCA. **(G)** Heatmap of the simulated RACIPE models for the best CGC. (**H**) Scatterplot of the simulated gene expression data of original circuit projected onto the first two principal component (PC) axes of the expanded network’s simulated data. RACIPE models belonging to the same cluster are colored with the same color. **(I)** Scatterplot of the simulated data of the expanded network projected onto its first two PCs. **(J)** Scatterplot of the simulated data of the best CGC projected onto the first two PCs of the simulated data of the expanded network. (**K**) Boxplots of the distributions of the last scores (log-scale) obtained by each MCMC method for each network perturbation level. (**L)** Boxplots for the distributions of edge mismatches of the last-iteration circuits and the original circuit for each MCMC method and each network perturbation level.

Next, we compared the performance of circuit optimization using three different MCMC sampling methods: Metropolis-Hastings (MH), Simulated Annealing (SA) and Parallel Tempering (TE). f **Fig.2KL** summarizes the performance of all three MCMC methods for networks perturbed by different edge-perturbation levels. **Fig.2K** shows the last-iteration scores of the sampling methods for different levels of edge-perturbation. The score measures the mismatch between the expression profiles of the CGC and those of the large network -- the smaller the score, the better the matching. As expected, for each perturbation level the median TE score is lower than the median SA score, which is lower than the median MH score. It is remarkable that, for each MCMC method, the scores do not increase significantly as we increased the perturbation level from 12% to 50%. Only for the 60% edge-perturbation level, the median score of MH is slightly higher than the other median MH scores. Even for this high perturbation level, we can obtain low scores. The best overall score was obtained by a SA run from a network with 50% of edge perturbations. For each MCMC sampling method, we calculated the number of edge mismatches (existent *vs*. nonexistent, or different edge types) between the last-interaction CGC and the original circuit. For instance, the number of edge mismatches between the original circuit (**Fig. 2A**) and the optimal circuit (**Fig. 2D**) is three, because the latter has three edges not existing in the original circuit. As shown in **Fig.2L**, the number of edge mismatches is on average between four and six for all MCMC methods and perturbation levels. Interestingly, many of these mismatches are self-activation loops due to the mutual activations between genes from the same group in the expanded networks; but these self-activation loops are absent in the original circuit. Otherwise, we observed low levels of mismatches in all sampling methods. Overall, we found that the circuit optimization procedures are effective in identifying CGCs even with high levels of perturbations.

The second test case is a repressilator circuit(Elowitz and Leibler, 2000), consisting of three self-activating nodes *A, B*, and *C*, where *A* inhibits *B, B* inhibits *C*, and *C* inhibits *A* (diagram in **Fig.3A**). To generate the expanded network, we expanded each node to threes genes, *i*.*e*., *A* is expanded to genes 1, 2, and 3; *B* to genes 4, 5, and 6; *C* to genes 7, 8, and 9 (**Fig.3B**). Similar to the previous case, we expanded the circuit to a large GRN by connecting multiple copies of the original circuit with consistent edges. We also generated large networks with different levels of edge perturbations. **Fig.3CD** show an example of such a perturbed network with 50% of edge perturbations and the last-iteration CGC obtained from SacoGraci. In this example, the original circuit, the expanded network, and the CGC all generate very similar gene expression profiles, as illustrated in **Fig.3E-J**. Note that a repressilator is usually modeled to generate oscillatory dynamics (Elowitz and Leibler, 2000). However, from an ensemble of models with randomly generated kinetic parameters, the steady-state gene expression profiles capture the network states along an oscillatory trajectory(Katebi et al., 2020). This explains why the coarse-graining scheme works very well for an oscillatory system here, where we match the steady-state gene expression profiles. The last-iteration scores of all MCMC methods (**Fig.3K**) are mostly very low, indicating that all methods obtained CGCs with a very good fit most time. In fact, 90% of the MCMC runs produce the same last-iteration circuit as the original circuit. The original circuit also produces the lowest score in this case. The number of edge mismatches between the last-iteration circuits and the original circuit (**Fig.3L**) are much lower than those from the first example. We observed a slightly worse performance of circuit optimization for networks with high perturbation levels; yet TE usually has the most stable and best performance for those cases.

**Figure 3.**
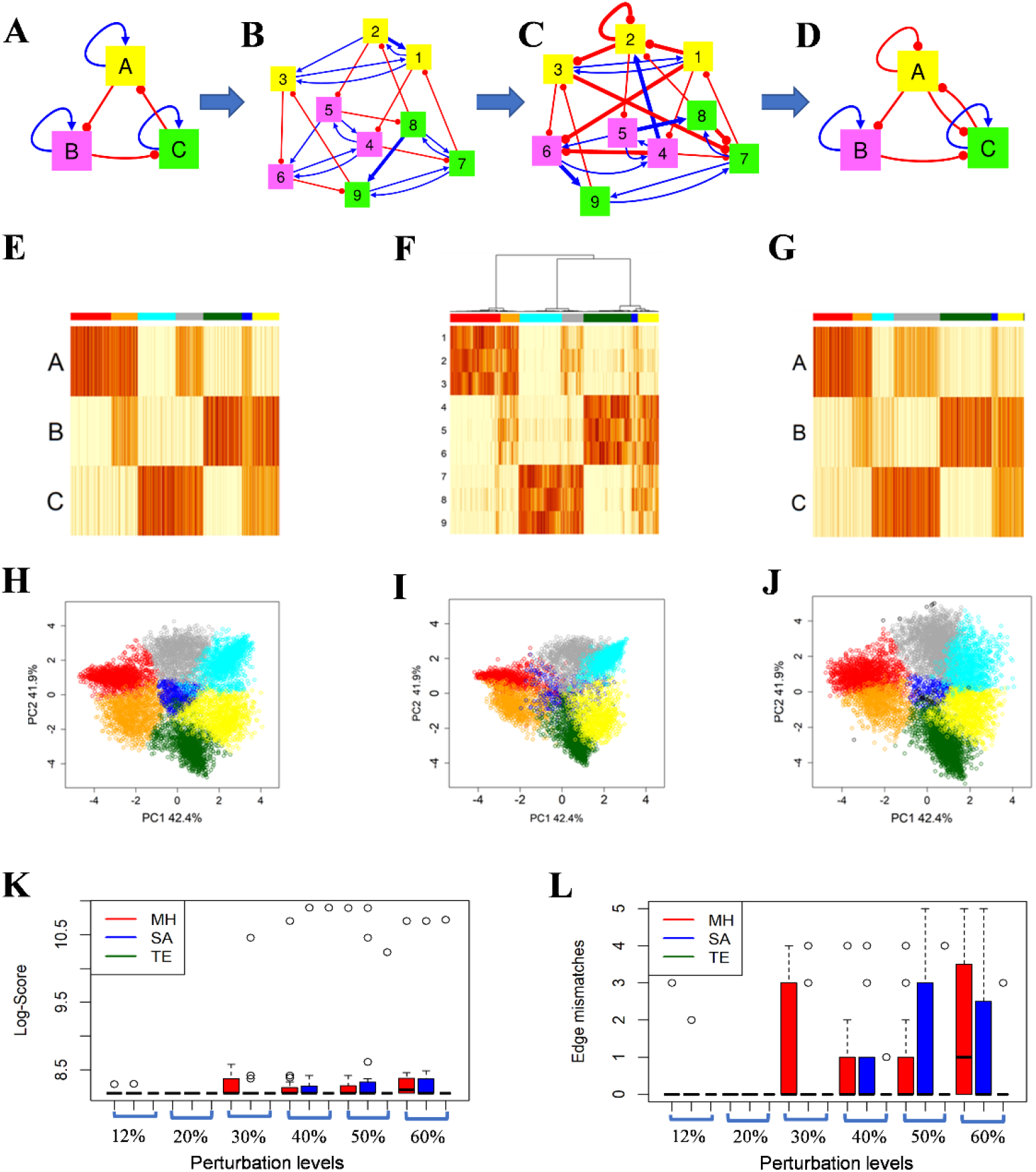
Benchmark tests on a synthetic gene circuit of a repressilator with self-activation loops. **(A)** Topology of the original circuit. **(B)** Topology of an expanded network. Every node of the original circuit is expanded into three nodes of the same color. Edges that will not be present in the perturbed network are shown in bold. **(C)** Topology of a perturbed network with a 50% edge perturbation level. Edges that have been added to the expanded network or have changed the sign are shown in bold. **(D)** Topology of the second best last-iteration CGC obtained after running all MCMC methods across all perturbation levels considered. The best last-score CGC is the same as the original circuit. (**E**) Heatmap of the RACIPE-simulated data for the original circuit. **(F)** Heatmap of simulated gene expression of the expanded network. It shows the hierarchical clustering analysis (HCA) of the RACIPE models using Ward.D as linkage method and 1 minus Pearson correlation as distance. Seven well-separated clusters are determined based on HCA. Gene clustering puts the genes of the same color into the same group. **(G)** Heatmap of the simulated RACIPE models for the second best CGC. (**H**) Scatterplot of the gene expression data of the original circuit projected onto the first two PCs of the expanded network’s simulated data. RACIPE models belonging to the same cluster are colored with the same color. **(I)** Scatterplot of the simulated data of the expanded network projected onto the first two PCs. **(J)** Scatterplot of the simulated data of best CGC projected onto first two PCs of the simulated data of the expanded network. (**K**) Boxplots of the distributions of the last scores (log-scale) obtained by each MCMC method for each network perturbation level. (**L)** Boxplots for the distributions of edge mismatches of the last-iteration circuits and the original circuit for each MCMC method and each network perturbation level.

### Coarse-graining a GRN of Epithelial-to-Mesenchymal Transition (EMT)

In the previous section, we showed the performance of SacoGraci on a series of large GRNs expanded from two synthetic circuits. Next, we illustrate the application of SacoGraci on four biological GRNs, including three literature-derived networks and a network generated by bioinformatics analysis.

The first example is a GRN responsible for the decision making of EMT. During EMT, epithelial cells undergo a phenotypic transition to mesenchymal cells by losing cell adhesion and gaining high motility (Nieto et al., 2016). EMT has been found to play crucial roles in embryonic development, wound healing, and cancer metastasis (Gupta and Massagué, 2006; Thiery et al., 2009). The EMT GRN, which consists of 13 TFs, 9 microRNAs, and 82 regulatory links between them (network topology shown in **Fig.4A**), was previously constructed (Huang et al., 2017) using the gene regulatory data from the literature and Ingenuity Pathway Analysis (Krämer et al., 2014).

**Figure 4.**
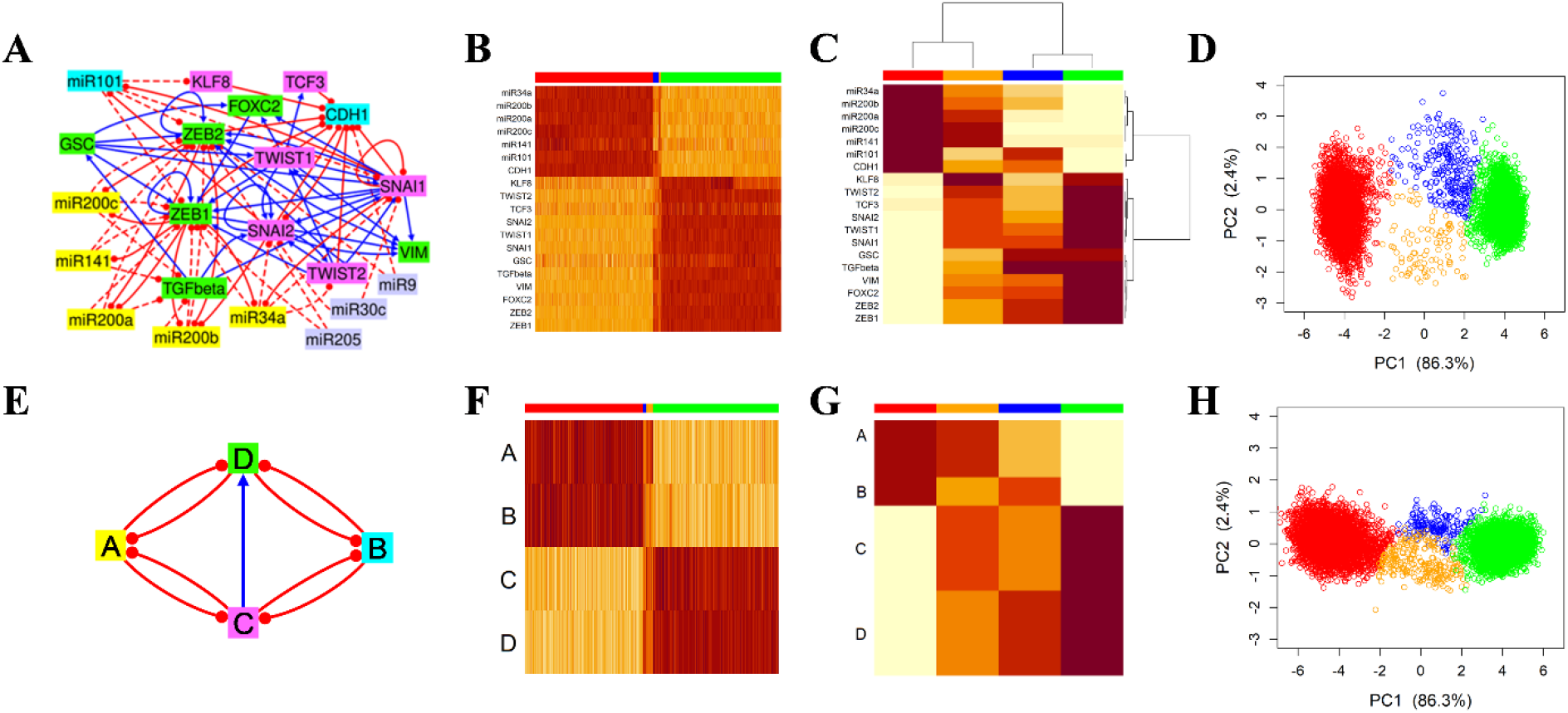
Coarse graining a gene network regulating epithelial-mesenchymal transition (EMT). **(A)** EMT network topology **(B)** Heatmap of the RACIPE-simulated gene expression profiles from the EMT network. Each row represents a network gene, and each column represents a simulated model. Clustering of the simulated RACIPE models was done using k-means clustering (k=4) on the 2D points obtained from the simulated data projected onto its first two PCs. **(C)** Heatmap of the median values of the simulated models for each gene in each cluster. Red and green clusters correspond to epithelial and mesenchymal states, respectively. Orange and blue clusters correspond to the hybrid states. Gene clustering was performed by HCA of this heatmap, where four gene clusters were identified. Genes belonging to the same gene cluster are illustrated with the same color as those in the network diagram shown in panel **A. (D)** Scatterplot of the simulated data of the full network projected onto its first two PCs. **(E)** Network topology of the optimized CGC (second best). **(F)** Heatmap of the RACIPE-simulated models from the optimized CGC. Each row represents a CGC node, and each column represents a RACIPE model. **(G)** Heatmap of the median expression values of the simulated data of the optimized CGC. Each row represents a CGC node, and each column represents a model cluster. **(H)** Scatterplot of the simulated data of the CGC projected onto the first two PCs of the full network (panel **D**). Models corresponding to the same cluster are illustrated with the same color.

We first applied RACIPE to the EMT GRN and obtained stable steady states from 10,000 models (one steady state per model). From hierarchical clustering analysis of the simulated gene expression profiles (**Fig.S1**), we observed two major clusters of models, representing the epithelial (E, with high CDH1 levels) and mesenchymal (M, with high VIM levels) states. We also observed a small fraction of models with intermediate levels of gene expression. However, the models do not form distinct clusters in HCA, thus they are difficult to be separated from the E and M states. But when the same data was projected to its first two PCs (**Fig. 4D**), we can clearly observe these models (colored in blue and orange) away from the E (red) and M (green) states. We performed k-means clustering (k = 4) of the data projected to the first two PCs and successfully identified four distinct clusters of models associated with the E, M states and two hybrid EMT states (EM I and EM II). We opted to do the clustering in a supervised manner, by providing the center clusters as the input to the 4-means algorithm. The model clustering is consistent with experimental observations of hybrid EMT phenotypes (Karacosta et al., 2019; Qiu et al., 2022; Zhang et al., 2014).

Once the model clustering has been determined, we then grouped genes based on the gene expression patterns of different EMT states. We started from the RACIPE-simulated gene expression profiles (**Fig.4B**), computed the median expression values for each gene in each model cluster, and then performed HCA again (**Fig.4C**). Here, the median values were computed in **Fig.4C** to emphasize on the gene expression differences in the rare hybrid EMT states. From the last step, we obtained the gene clustering, which we used to determine the nodes of CGCs. We decided that four gene clusters will be sufficient to capture the expression patterns. The gene grouping, as annotated in **Fig.4A**, puts miR34a, miR141, miR200a, miR200b and miR200c in node *A*; miR101 and CDH1 in node *B*; KLF8, TCF3, SNAI1, SNAI2, TWIST1 and TWIST2 in node *C*; and GSC, FOXC2, TGFβ, VIM, ZEB1 and ZEB2 in node *D*. As the genes miR205, miR9 and miR30c act as input nodes, *i*.*e*., nodes not regulated by other genes, they were not considered for gene grouping. Genes in node *A* are major miRNAs responsible for inducing the epithelial state (Zhang and Ma, 2012). CDH1 (*i*.*e*., E-Cadherin) in node *B* is known to be regulated by major EMT master regulators, such as SNAIL and ZEB families, and serves as a readout in previous EMT gene regulatory circuit models (Lu et al., 2013; Tian et al., 2013). MiR101 forms a toggle switch with SNAI1, while CDH1 forms a toggle switch with SNAI2. Also, miR101 inhibits ZEB1 and ZEB2, which are repressors for CDH1. All these interactions lead to a 0.91 Pearson correlation between CDH1 and miR101 in simulated gene expression. Thus, CDH1 and miR101 were grouped together in node *B*. Genes in nodes *C* and *D* are mainly regulators responsible for mesenchymal states.

With the model clustering and gene grouping being determined, SacoGraci can then be applied to identify the optimal CGCs with MCMC sampling. We ran all three MCMC methods and searched for the best performing CGCs. An example of such CGC (second best last-iteration circuit) is illustrated in **Fig.4E**. As shown in the heatmaps (**Fig.4FG**) and PCA results (**Fig.4H**), the simulated gene expression profiles from the CGC resemble those from the original EMT network, indicating a successful coarse graining. While the symmetry of the CGC would suggest that nodes *A* and *B* play similar roles and they might as well be combined into a single node, we note that having both *A* and *B* helps to distinguish between the two hybrid states.

In the optimal CGC, node *A* forms two double-negative feedback loops with nodes *C* and *D*, respectively. These circuit motifs are consistent with the double-negative feedback loops formed by the miR34 and SNAIL families (as in the case of the *A-C* loop) and those formed by the miR200 and ZEB families (as in the case of the *A-D* loop). Both SNAIL and ZEB inhibit CDH1, CDH1 inhibits SNAI2, and miR101 inhibits ZEB; all of these explain the *B-C* and *B-D* double-negative feedback loops. Interestingly, the CGC has a directed interaction from node *C* to *D*, consistent with the gene regulation from SNAIL to ZEB. It is also supported by the biological observations that the node *C* genes, such as TWIST and SNAIL families, are usually activated prior to the activation of ZEB family (Jia et al., 2017; Lu et al., 2013; Zhang et al., 2014). Moreover, the mono-directional interaction is also largely consistent with the edge type distributions from the large GRN. Taken together, the CGC captures the major regulatory features of the large EMT network. Note that in another CGC model (best last-iteration CGC, as shown in **Fig.S2**) *C-to-D* becomes bidirectional, but *C-to-A* becomes mono-directional. Presumably some asymmetry is needed to match well with full-network gene expression. Overall, the former CGC model is more consistent with experimental evidence of EMT.

### Coarse-graining a GRN of small cell lung cancer (SCLC)

The second example is a GRN of SCLC (Udyavar et al., 2017) consisting of 33 transcription factors (**Fig.5A**). Previous studies have shown that the network allows neuroendocrine/epithelial (NE), mesenchymal like (ML), and hybrid phenotypes (Udyavar et al., 2017). We also observed gene expression clusters associated with these phenotypes from the simulations of the SCLC GRN using RACIPE (**Fig.5BC**). From the HCA of the simulated gene expression data (**Fig.5B**), we decided to group the genes into four nodes (grouping scheme illustrated in **Fig.5A** by gene colors). With the defined model and gene grouping scheme, we performed the circuit optimization with MCMC meth, where the best last-iteration CGC is shown in **Fig.5D**.

**Figure 5.**
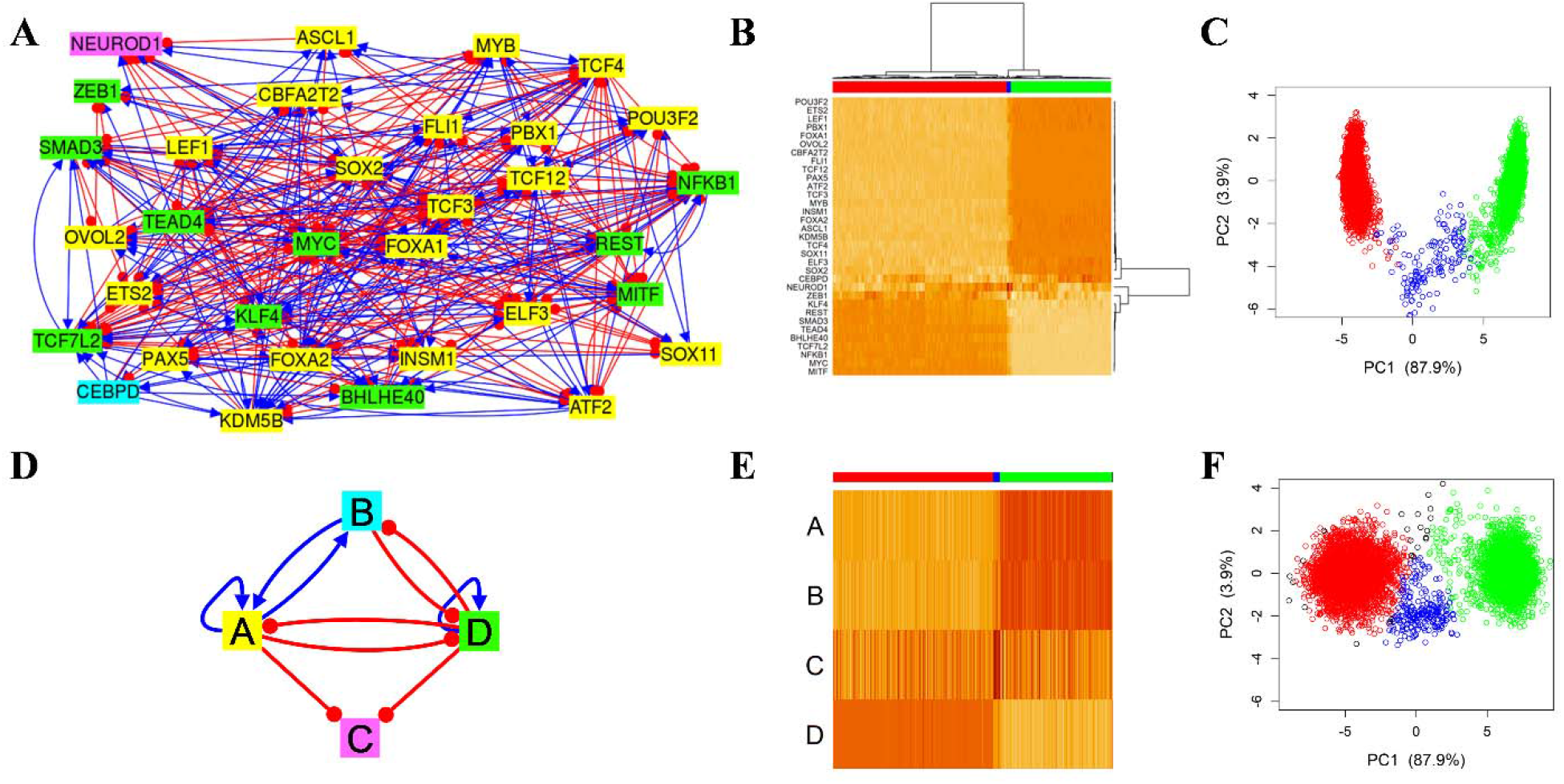
Coarse graining a gene network involved in small cell lung cancer (SCLC). **(A)** SCLC network topology **(B)** Heatmap of the RACIPE-simulated gene expression profiles from the SCLC network (HCA with Ward.D2 and Pearson correlation). Each row represents a network gene, and each column represents a simulated model. Three model clusters and four gene clusters were determined based on this clustering. Genes belonging to the same gene cluster have the same color in the network diagram shown in panel **A. (C)** Scatterplot of the simulated data of the SCLC network projected onto its first two PCs. (**D**) Network topology of the best CGC. **(E)** Heatmap of the simulated data of the best CGC. Each row represents a CGC node, and each column represents a simulated model. **(F)** Scatterplot of the simulated data of the best CGC projected onto the first two PCs of simulated data of the full network. Models corresponding to the same cluster are illustrated with the same color. There is a very small fraction (∼0.7%) of models unclassified to any cluster (black points in panels **E** and **F**).

In the optimal CGC, node *A* consists of ASCL1, a SCLC biomarker and a transcription factor involved in neuronal commitment, and some other genes involved in neuronal development and differentiation, such as SOX2, POU3F2 or TCF3 (Graham et al., 2003; Gribble et al., 2009; Kasher et al., 2016). Node *B* consists of one single gene CEBPD, which has diverse cellular functions depending on biological contexts. For example, CEBPD can act as a tumor repressor in pancreatic ductal adenocarcinoma (Hartl et al., 2020) or can contribute to proinflammation when activated by IL-6 (Cantwell et al., 1998). Node *C* also consists of one single gene NEUROD1, another SCLC biomarker associated with cell migration and metastasis (Ikematsu et al., 2020). Node *D* consists of mainly mesenchymal biomarkers, like ZEB1, and oncogenes, like MYC and TCF7L2. The topology of the CGC (**Fig.5D**) reveals a toggle-switch like topology with an added readout node (node *C*). Nodes *A* and *B* stimulate each other, and both form double-negative feedback loops with node *D*. Note that the nodes like *B* and *C* contain a single gene, which may suggest the importance of these genes in determining the behavior of the GRN. The optimal CGC is slightly larger than a previous reduced model (Chauhan et al., 2021) which combines the genes from *A* and *B* into one single node. When we applied SacoGraci to the reduced (three-node) model we obtained slightly worse results. The best three-node CGC (**Fig.S3C**, green cluster has a median PC1 value of 5.61) did not capture the epithelial state in the full network (green cluster, with median PC1 value of 7.20 in **Fig.5C**) as well as the best four-node CGC (**Fig.5F**, green cluster has a median PC1 value of 6.64). Moreover, we modeled CEBPD as a separated node because of its distinct gene expression, as seen in the full GRN (**Fig.5B**).

### Coarse-graining a GRN of gonadal sex determination (GSD) in human

The third example is a GRN established to model the differentiation process of GSD (Ríos et al., 2015). GSD is an important process during sex development, where bipotential gonadal primordium (BGP) differentiates either into testes or ovaries (Yang et al., 2019). Mutations in genes involved in GSD regulation can lead to disorders in sex development (Ohnesorg et al., 2014). From the simulation of 10,000 RACIPE models of the GSD GRN, we identified six clusters of gene expression states (**Fig.6B-D**), which can be associated with various cellular states during GSD. In particular, the red and green clusters correspond to the differentiated Granulosa cells and Sertoli cells, respectively. The purple and orange clusters correspond to undifferentiated precursor cells, with the orange one closer to the Granulosa lineage and the purple one closer to the Sertoli lineage. The major difference between the differentiated and undifferentiated states is the levels of UGR, the node in the GRN consisting of LHX1, LHX9, EMX2, PAX2 and PAX8 genes and representing the urogenital ridge, an embryonic structure precursor of the gonads. UGR gene expression indicates whether the differentiation process takes place (high UGR levels) or not (low UGR levels) (Yang et al., 2019). The blue cluster has relatively high UGR, GATA4, and Granulosa cell-specific genes, such as WNT4 and FOXL2; thus, it likely corresponds to Granulosa cells, but not fully differentiated. The yellow cluster has low levels of UGR and high levels of Sertoli cell-specific genes, likely associated with a genetic disorder due to the low level of GATA4 (Lourenço et al., 2011). Note that the RACIPE modeling was able to identify much richer cellular phenotypes than the previous Boolean network modeling.

**Figure 6.**
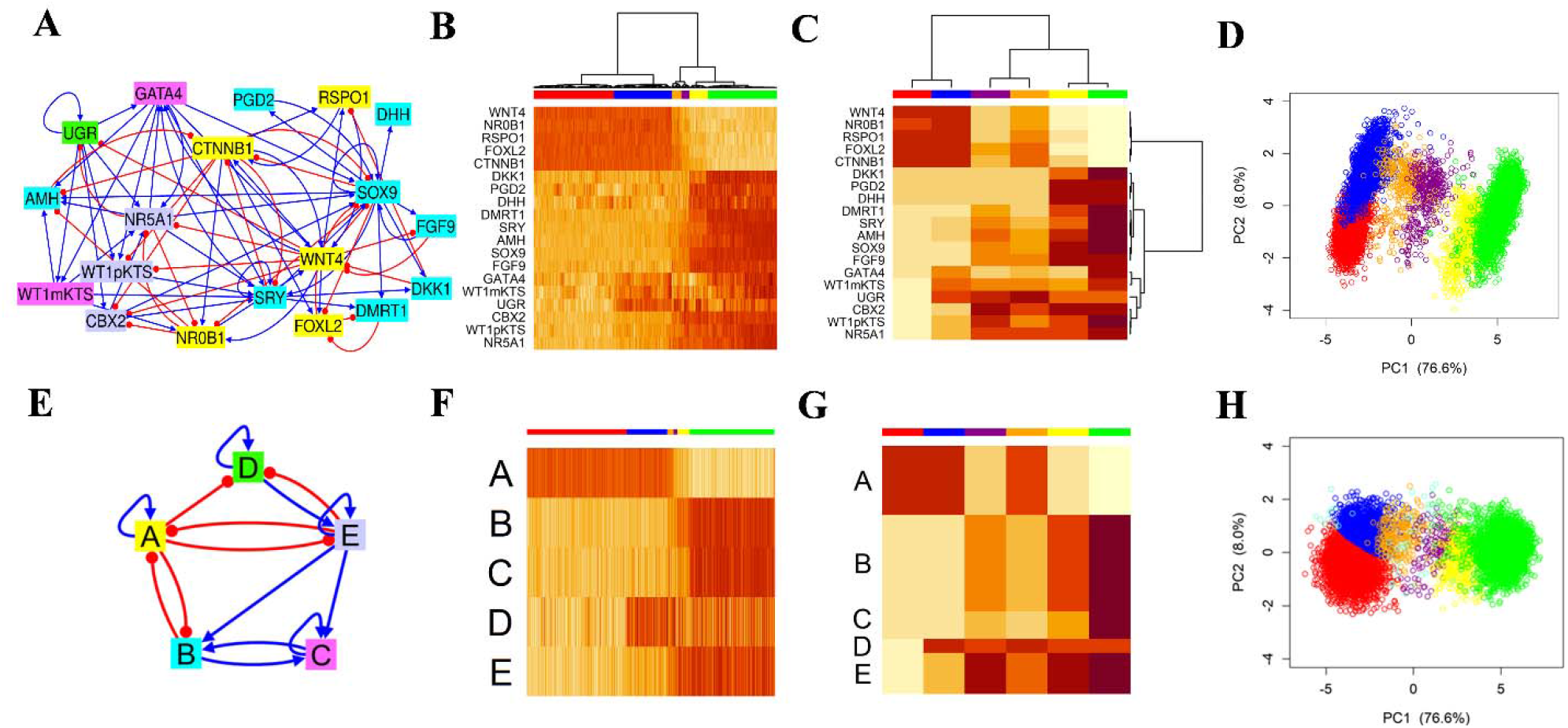
Coarse graining a gene network in human gonadal sex determination (GSD). **(A)** GSD network topology **(B)** The heatmap of the simulated gene expression profiles of the GSD network, where six model clusters were determined. **(C)** The heatmap of median values of the simulated models (HCA with Ward.D2 and Pearson correlation). From the HCA, five gene clusters were also identified. **(D)** Scatterplot of the simulated data of the GSD network projected onto its first two PCs. (**E**) Network topology of the best CGC. **(F)** Heatmap of the simulated models of the best CGC. **(G)** Heatmap of median expression values of the simulated data of the best CGC. **(H)** Scatterplot of the simulated data of the best CGC projected onto the first two PCs of the simulated data of the full network. Models corresponding to the same cluster are illustrated with the same color. The aquamarine models in panels **F** and **H** correspond to models unclassified to any cluster.

Next, from the median gene expression profiles of each model cluster (**Fig.6C**), we identified five gene clusters from HCA (the grouping scheme illustrated with colors in **Fig.6A**) for coarse-graining. **Fig.6E** shows the topology of best last-iteration CGC from SacoGraci. Note that the CGC captures very well all the six states of the original GRN, even for those states with similar gene expression profiles, such as the two undifferentiated states (purple and orange). In the CGC, node *A* consists of Granulosa cell-specific genes such as FOXL2, CTNNB1, RSPO1 or WNT4. The other gene in the node, NROB1 (a marker gene of gonadal primordium BGP) is known to act antagonistically to SRY (Lalli and Sassone-Corsi, 2003; Swain et al., 1996), the gene that triggers Sertoli cell differentiation. Network begins differentiation toward ovaries when all the genes in node A are active. When active, CTNNB1 and FOXL2 inhibit Sertoli cell differentiation (Ríos et al., 2015). Node *B* contains genes specific to Sertoli cells. The Sertoli cell differentiation pathway gets activated by first activating SRY, which activates SOX9 and then the rest of the genes in *B* in an activation cascade. The SOX9 activation is also followed by the repression of the Granulosa cell differentiation pathway (Koopman, 1999; Ríos et al., 2015). Node *D* contains one single gene UGR, again confirming its important role in the GRN of GSD. Lastly, the genes from nodes *C* and *E* are specific to non-differentiated BGP.

The CGC topology indicates a toggle switch-like behavior with many possible intermediate states. There are two main states of the CGC, one with high levels of node *A* and the other with high levels of nodes *B, C* and *E*. These two states are established and maintained by (1) the toggle switch between nodes *A* and *B*; (2) the toggle switch between nodes *A* and *E*; (3) the activation links between nodes *B, C* and *E*; (4) node *D* that activates nodes *C* and *E* and is repressed by node *A*. But node *D* is repressed by node *E*, which could explain the existence of the undifferentiated states. Taken together, the optimal CGC not only captures the major cellular states of the GSD network, but also sheds light on the logics of gene regulation in the complex system.

### Coarse graining a GRN of TGF-β-induced EMT specific to a cancer cell line

Unlike previous examples where GRNs are literature-based, the fourth example is a GRN constructed by a combined bioinformatics and systems biology modeling approach (the diagram of the GRN topology illustrated in **Fig.7A**) (Ramirez et al., 2020) using time-series scRNA-seq data of TGFB1-induced EMT in ovarian cancer cell line OVCA420 (Cook and Vanderhyden, 2020). From RACIPE simulations, we identified three gene expression clusters that can be associated with the mesenchymal (M) state (green), and two epithelial-like states (the first one E I, in red, the second one, E II, in blue) (**Fig.7BC**). In this case, we found it is sufficient to capture gene expression variations with four gene groups, thus being selected as the grouping scheme for coarse graining.

**Figure 7.**
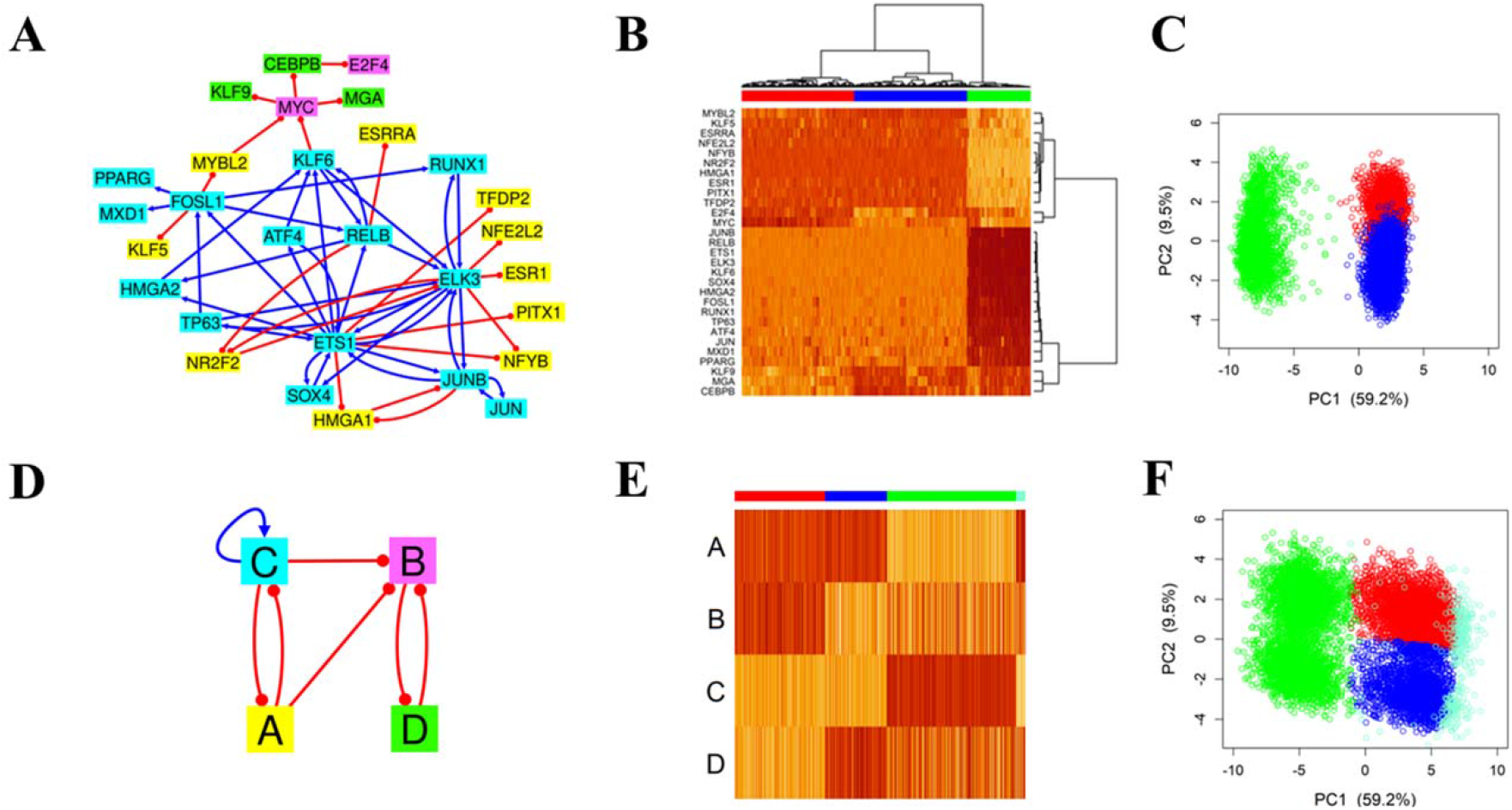
Coarse graining a network of TGFB1-induced EMT in OVCA420 cancer cell line. **(A)** OVCA420 network topology **(B)** The heatmap of the simulated gene expression profiles of the network (HCA with Ward.D2 and Pearson correlation). From the HCA, three model clusters and four gene clusters were determined. **(C)** Scatterplot of the simulated data of the full network projected onto its first two PCs. (**D**) Network topology of the best CGC. **(E)** Heatmap of the simulated models of the best CGC. **(F)** Scatterplot of the simulated data of the best CGC projected onto the first two PCs of the simulated data of the full network. There is a small fraction (∼3%) of models unclassified to any cluster (aquamarine points in panels **E** and **F**).

**Fig.7D** shows the topology of best CGC obtained after coarse graining the network using the MH algorithm as the sampling scheme. The sampling space of possible topologies was very small in this case, so we did not need to apply the more computationally intensive methods, *i*.*e*., SA and TE. The CGC topology captures the main features of the full network. In the full network, the genes from node *C*, JUNB and ELK3, form toggle switches with the genes from node *A*, HMG1 and NR2F2, respectively. This is reflected in the CGC by the toggle-switch between nodes *A* and *C*. Moreover, MYBL2 from node *A* and KFL6 from node *C* repress the MYC gene from node B, which explains the inhibitory edges from *A* to *B* and *C* to *B*, respectively. These interactions also explain how the E II state is allowed by the CGC.

Lastly, we also compared the performance of the three MCMC methods, MH, SA and TE, in the above-mentioned applications to four biological networks. **Fig.S4** shows the boxplots of last-iteration scores obtained by these methods for the four networks. Particularly for the third example where five-node CGCs were optimized (**Fig.S4C**), we observed that on average TE method generates CGCs with lower scores than those from MH and SA, presumably because of larger sampling space allowed by the TE method. When modeling circuits with less than four nodes, however, SacoGraci has similar performance among MH, SA and TE (**Fig.S4ABD**). Note that all these biological GRNs contain a high level of redundancy, which explains why the performance of network coarse graining is consistently high.

## Methods

### Simulating GRNs with RACIPE

We applied an ordinary differential equation (ODE)-based method, namely random circuit perturbation RACIPE, implemented as an R package sRACIPE (Kohar and Lu, 2018), to simulate the stable steady states of a GRN. RACIPE takes a network topology as the input and builds a system of ODEs that describes the network’s dynamics. By randomly sampling the kinetic parameters of the ODE system from biologically relevant ranges, RACIPE builds an ensemble of models, each of which as an instantiation of ODEs where all kinetic parameters and initial conditions are randomly specified. These kinetic parameters reflect different characteristics of the network, such as production and degradation rates for genes and parameters from shifted Hill function for regulatory interactions. See the original RACIPE study (Huang et al., 2017) for details of the methodology. Each model is simulated by an ODE solver until convergence to a stable steady state. Finally, we obtained a set of gene expression profiles from the analysis of an ensemble of models. A total of 10000 RACIPE models were typically generated to ensure we captured robust features of the GRN. The steady state gene expressions from these models usually form robust clusters, which can be further associated with distinct cellular states. Thus, we can predict the potential cellular states only from the network topology.

### Model clustering and gene grouping

From the RACIPE-simulated gene expression profiles of the large GRN, we aim to determine the distinct patterns of gene expression and distinct groups of genes with shared dynamical behavior in the network. To achieve this, SacoGraci performs hierarchical clustering analysis (HCA) to determine the number of model and gene clusters by default. The other method we used for model clustering was k-means clustering. However, users can also provide their own choices of the grouping scheme. For the HCA, we typically use a distance function of one minus Pearson correlation between the gene expression profiles of two models and the *Ward’s* D2 (Ward, 1963) minimum variance method as the linkage method. The only exception was for the case of the synthetic repressilator circuit with activation loops, where we used the older Ward’s D method.

To evaluate the quality of the grouping scheme, we also recommend to visually examine the heatmap of the clustered models and the scatter plot of gene expression profiles with low dimensional projection, using methods such as principal component analysis (PCA). With a high-quality grouping scheme, the clusters obtained from hierarchical clustering should form dense, compact scatterplot regions. Clusters that have similar expression patterns and have very close or overlapping scatterplot regions might be combined. For gene grouping, users can explore different number of gene clusters and/or incorporate prior knowledge (such as the number of gene families involved in the biological context) into the pipeline. Each of the gene group from the clustering analysis will be a node of the CGC. In our test cases we chose different number of gene clusters (*e*.*g*., from three to five) to explore CGCs of different sizes.

### Sampling coarse-grained circuits

In the previous step, SacoGraci defines the CG nodes from the clustering analysis. Now, we will determine how the edges of CGCs are sampled. For each ordered pair *(A, B)* of gene clusters, we determine the probability distribution of the edge type (activation, inhibition, no edge) from *A* to *B*. To do so, we examine all possible edges from genes in *A* to genes in *B* from the full network and calculate the proportion of activation and inhibitory edges. This will give us the probability of an activation edge from *A* to *B* and probability of an inhibitory edge from *A* to *B*, respectively. The remaining fraction determines the probability of having no edge from *A* to *B*.

As mentioned above, the nodes of a CGC correspond to gene groups of the full network. Then, to build a CGC we sample edges for all ordered pairs of nodes. The edge types are sampled according to the above calculated edge type distributions. The only restriction we impose here is to get a connected topology with no input nodes. RACIPE-simulated gene expression of an input gene/node looks like noise because nothing regulates the expression of that input gene. That is why, whenever a large network has input nodes, we delete them and apply SacoGraci to the remaining network. To obtain optimal CGCs, we defined the scoring function (see next section) and utilized three Markov Chain Monte Carlo (MCMC) sampling methods, i.e., Metropolis-Hastings (MH), Simulated Annealing (SA) and Parallel Tempering (TE) (Liu, 2008). Details about the implementation of MCMC methods are given in **SI text**.

### Scoring function for circuit optimization

A scoring function is defined to compare the simulated gene expression profiles of a CGC with those of a full network. We first determined the number of model clusters, *n*_C_, of the full network. Then each simulated model *i* of the CGC was expanded to a model in full network’s gene expression space. The expanded model was assigned to the closest cluster of the full network (*C*_i_) according to a normalized distance *E*_i_, defined as the ratio of the Euclidian distance from the expanded model to the cluster center and the cluster radius. See **SI text** for the definition of cluster center and radius. If the expanded CGC model is too far away from any full-network cluster (i.e., *E*_i_ < *1*), it would be regarded as an unassigned model. The total score is defined as

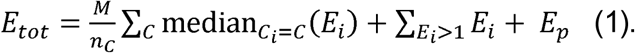

The first term evaluates the closeness of CGC simulated gene expression to full-network model clusters (*C*), where the summation is over all model clusters. *M = 10000* is the total number of CGC models. The second term gives penalties to the CGC models not assigned to any cluster. In addition, if for a certain cluster *C*, the number of assigned CG models is less than half of the full-network models in the same cluster *C*, then a penalty term of 50000 (*E*_p_) is added to *E*_tot_. Details about the scoring function are given in **SI text**.

### Expanding a small circuit and perturbing a full network

For benchmarking the MCMC sampling methods to obtain optimal CGCs, we started with a small gene circuit (the ground truth) and expanded it to a large network that reproduces the similar gene expression patterns of the circuit. To achieve this, we replaced each node in the circuit by three or four genes. We connected the genes corresponding to the same node by many activation links. That ensures that these genes will have a similar gene expression pattern. For two nodes connected by a directed edge in the small circuit, we connected the corresponding genes by 3 to 6 edges of the same type going in the same direction. Here, we created multiple copies of the original circuit (four copies for the coupled toggle-switches, three for the repressilator) and connected them with edges consistent with edges of the original circuit.

To evaluate the robustness of the coarse graining algorithm when some inconsistent interactions are presented in the expanded network, we generated networks with perturbed topologies. To perturb a network by a certain degree, we changed its topology as follows. For a certain perturbation level *a* and a network of *h* edges, we would need to make a total of *ah* edge changes, each of which is either change of the sign (activation or inhibition) of an edge (50% chance), adding an edge (25% chance), or deleting an edge (25% chance). We required that the resulting perturbed network to be connected and have no input nodes. The perturbation levels *a* were chosen from {12%, 20%, 30%, 40%, 50%, 60%}. To ensure a good coverage of the sampling space, for each perturbation level we first randomly generated 20 perturbed topologies. Then, for each one of them we calculated average number of edge mismatches with the other 19 topologies. We chose the top ten topologies with the most mismatches as our perturbed networks. This ensures that the chosen perturbed networks are most different from one another.

## Discussion

In this study, we introduced SacoGraci, a new coarse-graining algorithm for gene regulatory networks based on ensemble-based mathematical modeling, dimensionality reduction, and gene circuit optimization. The algorithm requires the topology of a GRN as the only input to produce the CGCs that are best at reproducing the gene expression profiles of the large GRN. We benchmarked the effectiveness and robustness of our method using two small synthetic circuits, each of which was expanded to large networks with different levels of edge perturbation. In both cases SacoGraci could successfully reproduce the gene expression profiles and recover the topology of the original synthetic circuit, even when the edge perturbation level was as high as 60%. Furthermore, we successfully applied network coarse-graining to several literature-based and bioinformatics-constructed GRNs. We expect SacoGraci to contribute to a better understanding of the gene regulatory mechanisms of biological networks. We also expect the algorithm to be developed into a generally applicable framework able to coarse grain many other types of biological networks.

From our benchmark tests and biological network applications, we observed the following strengths of our approach. First, in all cases we could reproduce GRN’s expression patterns using CGCs of small sizes. These circuits usually have only three to five nodes, but they are already sufficient in modeling large biological networks with up to six distinct cellular states. Second, the identified CGCs usually can preserve rare populations of cellular states. This is due to the big penalty term in the scoring function that disfavors the CGCs not able to capture all states. Third, the performance of the method was sometimes insensitive to the choice of number of coarse-grained nodes. For instance, in the case of the SCLC network, desired results have been obtained for either three-node circuit or four-node circuit (**Fig.S3** and **Fig.5**). Users may wish to rely on biological insights to select the most appropriate size of CGC. Fourth, the method worked well to handle large networks, *e*.*g*., those containing as many as 33 genes and 357 edges. The computational cost in general increases exponentially with the CGC size due to the increase in sampling space of CGC topologies. Restrictions imposed on what topologies can be sampled, *e*.*g*., due to edge type distributions derived from the large GRN (see Methods), can make the computational cost sub-exponential. For the most computationally intensive case we tested, *i*.*e*., coarse-graining GSD using five-node circuits, it took 4 hours of running time for MH and SA and 5-6 hours for TE (for two NVIDIA K80 GPU cards, with 2496 GPU cores per card). Here, the MH and SA methods usually converged after at most 400 iterations, while TE converged after around 200-250 iterations.

While SacoGraci proved very effective in finding CGCs that reproduce the behavior of a GRN to a great extent, there could be a few limitations in certain situations. First, in the current approach, we assumed the RACIPE-simulated gene expression profiles form clusters of spherical shape, which we use to approximate the model assignment to each gene expression state. While the current scoring function works very well in our test cases, this may generate assignment errors in some rare cases, where the spherical assumption is not satisfied. One potential solution is to define a new distance function between gene expression distribution of a GRN and that of a CGC using metrics, such as Kullback-Leibler divergence, Kolmogorov-Smirnov statistic test, or earth mover’s distance. Second, we chose the number of CG nodes based on the number of gene clusters that can reproduce the gene expression patterns of various model clusters. But there is no guarantee that, for instance, using four-node CGCs would get us much better results than using three-node CGCs. It would be an interesting question to identify the optimal number of coarse-grained nodes through optimization. Third, the quality of network coarse-graining and computational cost might strongly depend on the choice of the sampling method. Here, we used three MCMC methods to sample candidate CGCs. The most sophisticated method we applied, the parallel tempering algorithm, produced the best overall results, but was usually slowest for large GRNs with large sampling space. Some other sampling algorithms might further improve the combined speed and accuracy, such as genetic algorithms or some other heuristic methods, especially for cases where the number of coarse-grained nodes is more than five. For CGCs with less than five nodes, the simulated annealing method would suffice to perform well within a short amount of time. Lastly, the current scoring function was designed to match steady-state gene expression distributions, but not necessarily gene expression dynamics. It would be helpful to incorporate the information of gene expression time dynamics during the circuit optimization.

## Acknowledgments

This work is supported by the National Institute of General Medical Sciences of the National Institutes of Health under Award Number R35GM128717, and by startup funds from Northeastern University.

## Author contributions

*C. C. and M*.*L. designed the computational algorithm and wrote the manuscript. C*.*C. performed the research. M*.*L. conceived and supervised the project*.

## Data Requirements

The SacoGraci R package, tutorial script, and example codes are available at the following link: (https://github.com/lusystemsbio/SacoGraci)

