## Supplementary material for "A Data-Driven Optimization Method for Coarse-Graining Gene Regulatory Networks": SI

**SI Text**

**1. Defining the scoring function for circuit optimization**

A scoring function is defined to compare the simulated gene expression profiles of a CGC with those of a full network. For each CGC, we generate 10000 models with randomly selected kinetic parameters using RACIPE. Each model is simulated by an ODE solver until convergence to a stable steady state, which is represented by a vector of gene expression values:$x_{i}=\left( g_{i}^{1},g_{i}^{2},\ldots,g_{i}^{N} \right)$ , where $N$ is the number of genes in the network and $g_{i}^{j}$ is the expression value of $j^{th}$gene in the $i^{th}$ model.

For each model cluster $C$ of the full network, we define its center, $O_{C}$, as the medium of the gene expression vectors $x_{i}$ for every model $i$ in the cluster $C$. We prefer medium to other possible choices, such as centroid, as median of a group of values is more robust to outliers than other statistics. Next, we permute all 10,000 RACIPE-simulated gene expression vectors. The permuted gene expression vectors take the form

$x_{i,perm}=\left( g_{i_{1}}^{1},g_{i_{2}}^{2},\ldots,g_{i_{N}}^{N} \right)$ (1),

where $i_{1},i_{2},\ldots,i_{N}$ are values sampled from the set $\left\{ 1,2,\ldots,M \right\}$, and the total number of models is $M=10000$. Then, we define the radius of model cluster $C$, $r_{C}$, as the $\left[ \alpha*M \right]^{th}$ shortest Euclidian distance from any of the permuted gene expression vectors to the center $O_{C}$. Here [] is the integer part function and $\alpha$ is a parameter selected from the set $\left\{ 0.01,0.05,0.10,0.15,0.20 \right\}$ to control the cluster radius $r_{C}$. Parameter α varies from network to network but is the same for all model clusters corresponding to a given network. We also assume a spherical form of gene expression profiles for each model cluster. If $r_{C}$ is too small, many of the models initially belonging to cluster $C$ could be left outside of the sphere of radius $r_{C}$. If $r_{C}$ is too big, the sphere could include models with more than one gene expression pattern. In general, it is recommended to choose a rather small $r_{∁}$. For a given network, to choose α, we start with $\alpha=0.01$ and find the radius of each cluster. We check how many models were left outside of the spheres. We choose the next value of α if the number of left out models is bigger than 3% of $M$. The maximum possible value of α is 0.20.

With $O_{C}$ and $r_{C}$ computed for every model cluster of the full network, we then define the score of a given CGC as follows. First, an ensemble of $M$ models are simulated with RACIPE for the CGC. Second, the gene expression vectors of the CGC are expanded to the size of the full network by assigning the gene expression value of a CG node $A$ to all the genes belonging to $A$ in the full network. For instance, for a full network of 7 genes and a CGC containing 3 nodes $A,B,C$, if $A$ is a group of genes $A=\left\{ g^{1},g^{2},g^{3} \right\}$ and $B=\left\{ g^{4},g^{5} \right\}$ and $C=\left\{ g^{6},g^{7} \right\}$, then a gene expression vector of a model of CGC, $x_{i}^{CGC}=\left( g_{i}^{A},g_{i}^{B},g_{i}^{C} \right)$ can be expanded to a vector of the full network $x_{i}^{exp}=\left( g_{i}^{A},g_{i}^{A},g_{i}^{A},g_{i}^{B},g_{i}^{B},g_{i}^{C},g_{i}^{C} \right)$. Third, for each expanded vector $x_{i}^{exp}$ obtained in the previous step, we assign a scoring term $E_{i}$ as the minimum ratio of the Euclidian distance from $x_{i}^{exp}$ to the cluster center $O_{C}$ and the cluster radius $r_{C}$:

$E_{i}=\min_{C} \left\{ \frac{d(x_{i}^{exp},O_{C})}{r_{C}} \right\}$ (2).

When and only when $E_{i}$ is less than 1, we assign the model $i$ to the cluster that realizes the minimum:

$C_{i}=\underset{C}{\mathrm{argmin}} \left\{ \frac{d(x_{i}^{exp},O_{C})}{r_{C}} \right\}$ (3).

Sometimes, the ratio is less than 1 for multiple model clusters from the full network, indicating the expanded CG model is within the ranges of multiple model clusters. But according to Equation (3), the cluster with the minimum ratio is assigned. Eventually, each expanded CG model has an assigned score, and many of them are also assigned to the model clusters of the full network. For each model cluster $C$, we calculate the median score

$E_{C}=\underset{C_{i}=C}{\mathrm{median}} \left( E_{i} \right)$	(4).

Here, $E_{C}$ reflects how well models from a CGC represent the gene expression patterns of a given cluster $C$ of the full network. Fourth, we define the total score as

$E_{tot}=\frac{M}{n_{C}}\sum_{C} E_{C}+\sum_{E_{i}>1} E_{i}+E_{p}$	(5),

where $n_{C}$ is the number of model clusters of the full network. The second term in Equation (5) gives penalties to the expanded CG models not assigned to any cluster. In addition, if, for a certain cluster $C$, the number of assigned CG models is less than half of the full-network models in the same cluster $C$, then a penalty term of 50000 ($E_{p}$) is added to $E_{tot}$.

**2. Detailed implementation of Markov Chain Monte Carlo (MCMC) sampling methods**

We used three Markov Chain Monte Carlo (MCMC) sampling methods to obtain optimal CGCs: Metropolis-Hastings (MH), Simulated Annealing (SA) and Parallel Tempering (TE) (Liu, 2008). We assume the probability of a CGC having the score $E_{tot}$ is given by a Boltzmann distribution

$p(E_{tot}, T)\propto e^{-\frac{E_{tot}}{kT}}$,	(6)

where $T$ is the temperature, and $k$ is the Boltzmann’s constant (set to 1 here).

Since $E_{tot}$ only depends on the topology of the CGC, we use MCMC methods to sample different circuit topologies as follows. During every iteration, we only change an edge of the current CGC. An edge is picked at random from the current circuit and then we sample its type according to its edge type distribution determined by the topology of the full network.

Since the scoring function is based on RACIPE simulations of 10000 models, a new calculation of the score for the same circuit will produce a slightly different result. To improve the efficiency and robustness of the circuit optimization, we chose to calculate the score five times and take the average of the five scores as the final score, when we sample a CGC for the first time. If the circuit is sampled at a later iteration, its score is not calculated again.

For the synthetic circuit cases we chose ten different perturbed networks for each perturbation level (see “Expanding a small circuit and perturbing a full network” from the Method section). For each perturbed network, we run MH and SA twice and TE once as follows: 1) We chose two different coarse-grained topologies as the starting circuits for both MH and SA. 2) The same two circuits were also among the initial circuits of the TE method. The other initial circuits for the TE method were randomly generated according to the edge type distributions determined by the perturbed network. Total number of initial topologies used for TE was 24. 3) We run MH, SA and TE using the initial circuits determined above and the edge type distributions given by the perturbed network. All three MCMC methods were run for 1400 iterations

To coarse grain the EMT, SCLC and GSD networks we applied a similar simulation scheme. In each case we generated 20 CGCs that were used as starting circuits for both MH and SA. They were chosen in such a way that the number of edge mismatches between them were as big as possible. That was to ensure we cover different regions of the sampling space. The 20 initial circuits were also a part of the initial circuits of the TE simulations. Initial number of topologies for TE method was 24. Both MH and SA were run in parallel using 20 threads, each thread running a simulation. The whole TE procedure was run 10 times for each of these networks. We did not run SA and TE when we coarse-grained the OVAL420 network, as there were only a few CGC topologies to sample in this case. All three MCMC methods were run for 1400 iterations.

Before starting the MCMC methods, we run MH for 50 iterations. The purpose is to evaluate the magnitude of the scores of the CGCs. Knowing this we can decide on the temperature value we need to use for running MH and the temperature grids for SA and TE. We denote by $E_{MH}$ the last score we get from the preliminary MH run. It is recommended that the temperature value for MH, $T_{MH}$ satisfies the condition

$60<\frac{E_{MH}}{T_{MH}}<75$	(7).

This rationale of the criterion in Equation (7) can be understood as follows. If the score of the current circuit is $E_{cur}$, then the acceptance probability of a newly sampled circuit with score $E_{new}$is $e^{\frac{E_{cur}-E_{new}}{T_{MH}}}$. If we assume that $E_{cur}$ has the magnitude of $E_{EM}$, then, when the score is increased by 3%, the acceptance probability is larger than 10% for $T_{MH}$ satisfying the above constraint. This will ensure a sufficient acceptance of newly sampled circuits with slightly higher scores than the current one. In this study, for the coupled toggle switches example, we chose $T_{MH}=40$; for the other examples, we chose $T_{MH}=60$.

For SA method we developed a temperature schedule consisting of 14 decreasing values, with the method running 100 iterations for each temperature value. The first four temperature values ensured we did a deep exploration of the sampling space. Next three values were within the range around $T_{MH}$. As for reduction rule of the temperature, we chose two geometric rates, one for the first seven values and one for the last seven values. For the coupled toggle switches example, the starting temperature was 120, first decreasing rate was 0.85, and the second decreasing rate was 0.6. For the other examples, the starting temperature was 150, the first decreasing rate was 0.8, and the second decreasing rate was 0.6.

For TE method we created K=24 replicas of the original Monte Carlo Markov chain. Each replica has its own temperature and is engaged in its own Markov Chain Monte Carlo search for CGCs using MH. A temperature grid $T_{1}<T_{2}<\ldots<T_{K}$ is constructed with $T_{1}=1$corresponding to our target replica (our solution to the problem). The largest temperature was set to $T_{K}\cong\frac{E_{MH}}{30}$. That gives us a big chance of having an acceptance rate of at least 50% for replica with temperature $T_{K}$. Each replica can swap its state with one of the replicas of the neighboring temperature. The acceptance probability of swaps between replicas with temperatures $T_{i}$ and $T_{j}$ is

$\rho_{i,j}=min\left\{ 1,\frac{p(E_{tot,j},T_{i})p(E_{tot,i},T_{j})}{p(E_{tot,i},T_{i})p(E_{tot,j},T_{j})} \right\}$	(8),

where $p(E_{tot,i},T_{i})$ is the Boltzmann distribution with temperature $T_{i}$ and $E_{tot,i}$ is the score of the current circuit sampled by replica with temperature $T_{i}$.

To ensure a target swap rate for neighboring replicas around 0.4, we added new temperatures to the grid according to a robust feedback-optimized method (Hamze et al., 2010). First, we performed an update of sampled circuit topology for each replica using MH. Second, we proposed swaps between neighboring replicas L and L-1, L-1 and L-2, …,2 and 1, where L is the number of replicas. Third, we repeated the first and second steps $m$ times. Fourth, for each pair of neighboring replicas (*i, i-1*), the following quantity was calculated:

$\mathbb{Q}_{i,i-1}=\frac{1}{N_{swap}^{i,i-1}}\sum_{l=1}^{N_{swap}^{(i,i-1)}} ln\left( \rho_{i,i-1}^{l} \right)$	(9),

$N_{swap}^{\left( i,i-1 \right)}=m$ is the number of proposed swaps between replicas *i* and *i-1*, and $\rho_{i,i-1}^{l}$ is the acceptance probability for swapping *i* and *i-1* at the *l^th^* attempt. If $R_{i,i-1}=\left[ \sqrt{\frac{\mathbb{Q}_{i,i-1}}{ln(0.4)}} \right]>0$, then we add to the grid $R_{i,i-1}$temperatures, evenly spaced between $T_{i-1}$ and $T_{i}$_._ This temperature grid addition process was performed three times to make sure there were enough temperatures on the grid to prevent isolation of replicas. The values of $m$ for the three runs of the procedure were 50,100 and 150, respectively. After the temperature addition process, the first and second steps described above were performed for another 1100 times.

The initial temperature grid for the coupled toggle switches example was {1, 1.05, 1.1, 1.15, 1.2, 1.25, 1.3, 1.5, 2.0, 2.5, 3.0, 3.5, 4.0, 6, 9, 11, 13, 20, 28, 40, 55, 70, 85, 100}. For the other examples the initial grid was {1, 1.05, 1.1, 1.15, 1.2, 1.25, 1.3, 1.5, 2.0, 2.5, 3.0, 3.5, 4.0, 6, 9, 11, 13, 20, 28, 40, 55, 70, 90, 120}.

**SI Figures**

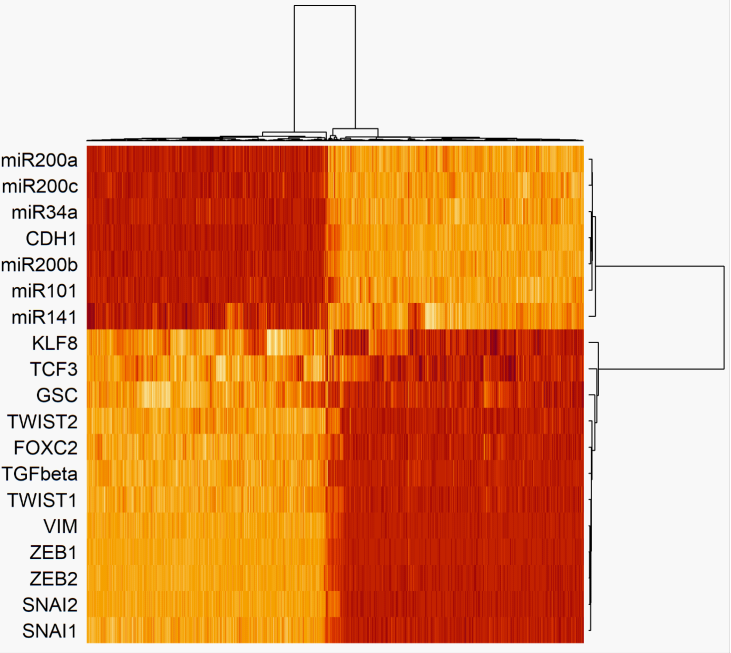

**Figure S1. Heatmap of the simulated RACIPE models for the EMT network.** Hierarchical clustering analysis was performed on the simulated RACIPE gene expression using Ward.D2 as the clustering method and one minus Pearson correlation as the distance function. Each row represents a gene, each column represents a RACIPE model.

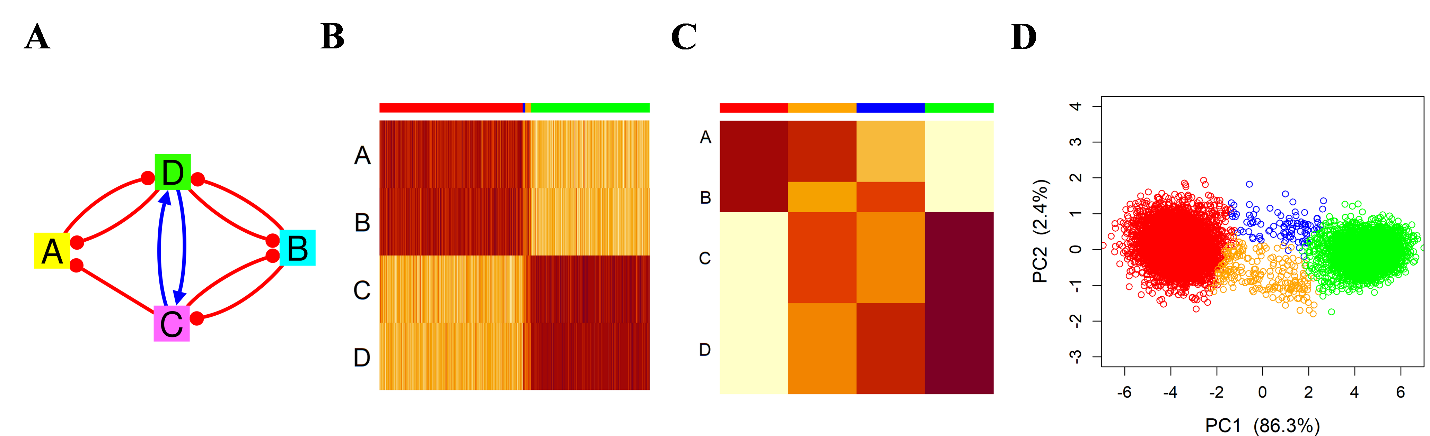

**Figure S2. Results of the best last-iteration CGC when coarse-graining the EMT network.** (**A**) Illustration of the CGC topology. **(B)** Heatmap of the RACIPE-simulated models corresponding to the best CGC. Each row represents a CGC node, and each column represents a RACIPE model. **(C)** Heatmap of the median expression values of the RACIPE-simulated data for the best CGC. Each row represents a CGC node, and each column represents a model cluster. **(D)** The embedded gene expression profiles for the best CGC projected onto the first two PCs of the full network’s simulated data. Models corresponding to the same cluster are shown with the same color.

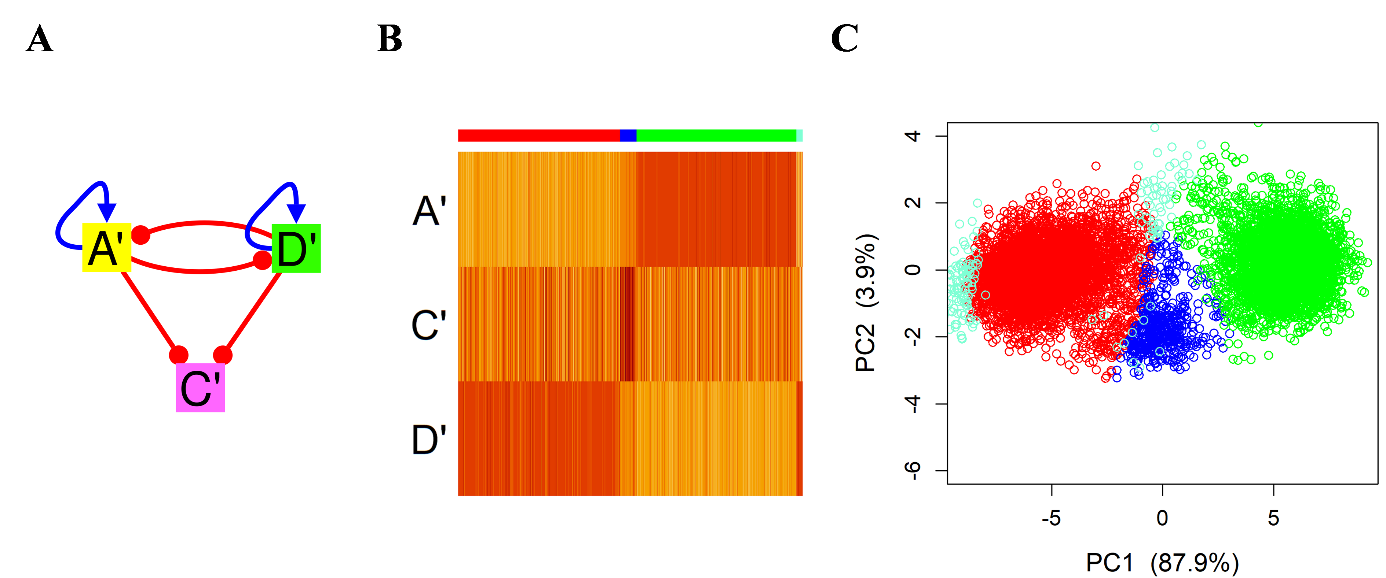

**Figure S3. Results of the best last-iteration CGC when coarse-graining the SCLC network using three-node circuits.** (**A**) Illustration of the CGC topology. The nodes A and B from the four-node gene grouping were combined into a single node A’, the nodes C and D were left unchanged; we renamed the nodes C and D from four-node gene grouping by C’ and D’, respectively. **(B)** Heatmap of the RACIPE-simulated models corresponding to best three-node CGC. Each row represents a CGC node, and each column represents a RACIPE model. **(C)** The embedded gene expression profiles for the best CGC projected onto the first two PCs of the full network’s simulated data. Models corresponding to the same cluster are shown with the same color. There is a very small fraction (~1.87%) of models unclassified to any cluster (aquamarine points in **B** and **C**).

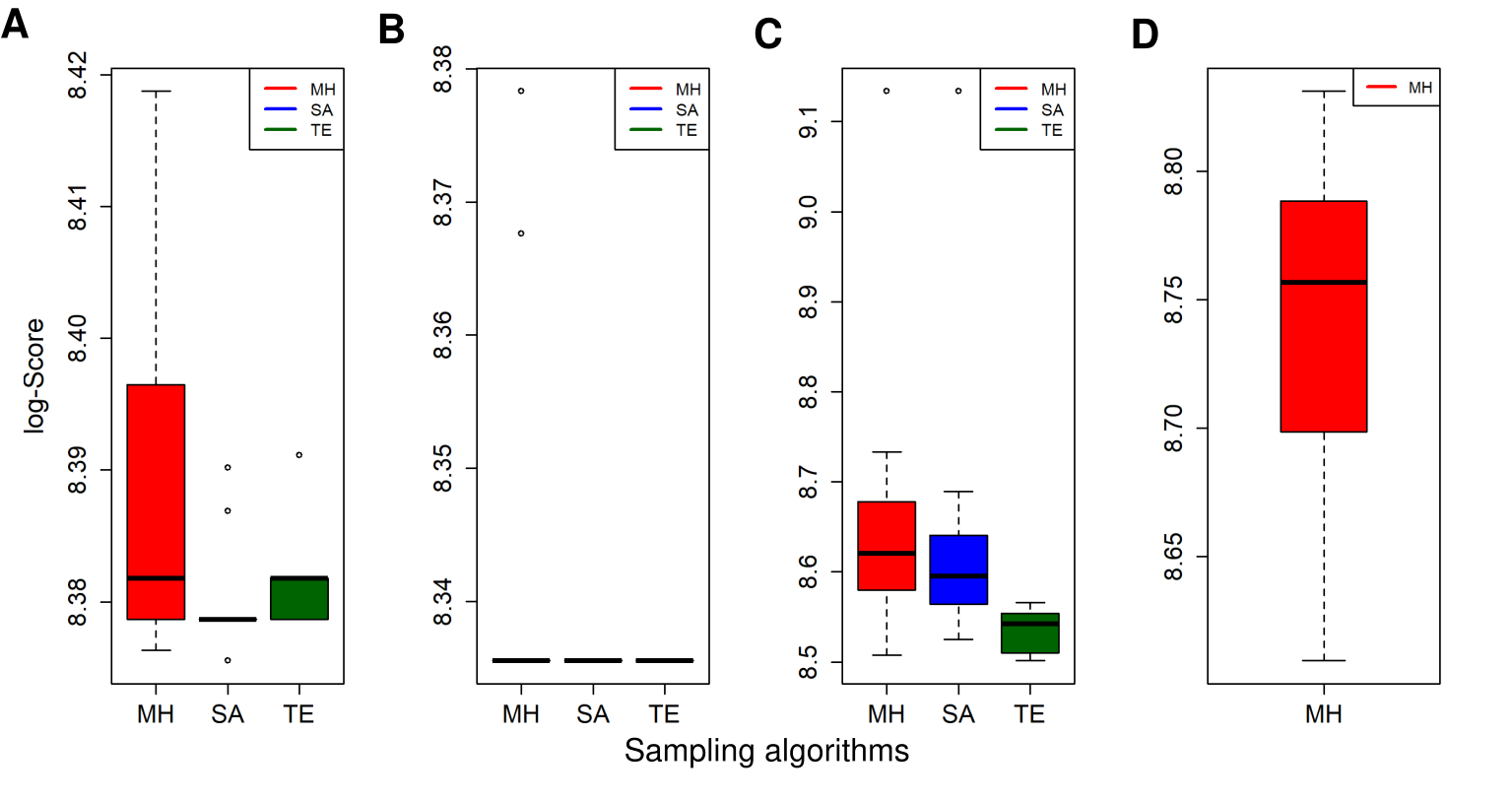

**Figure S4. Last-iteration scores obtained by each MCMC method. (A)** Boxplots of scores obtained when coarse-graining the EMT network. **(B)** Boxplots of scores obtained when coarse-graining the SCLC network. **(C)** Boxplots of scores obtained when coarse-graining the GSD network. **(D)** Boxplot of MH scores obtained when coarse-graining the OVCA420 network. Only MH is applied, as there are only a few circuit topologies to be sampled. The coarse-graining algorithm imposes strong restrictions on the sampling space in this case.
